# Backstepping mechanism of kinesin-1

**DOI:** 10.1101/838433

**Authors:** Algirdas Toleikis, Nicholas J. Carter, Robert A. Cross

## Abstract

Kinesin-1 is an ATP-driven molecular motor that transports cellular cargo along microtubules. At low loads, kinesin-1 almost always steps forwards, towards microtubule plus ends, but at higher loads, it can also step backwards. Backsteps are usually 8 nm, but can be larger. These larger backwards events of 16 nm, 24 nm or more are thought to be slips rather than steps, because they are too fast to consist of multiple, tightly-coupled 8 nm steps. Here we propose that not just these larger backsteps, but all kinesin-1 backsteps, are slips. We show firstly that kinesin waits before forward steps for less time than before backsteps and detachments; secondly that kinesin waits for the same amount of time before backsteps and detachments and thirdly that by varying the microtubule type we can change the ratio of backsteps to detachments, without affecting forward stepping. Our findings indicate that backsteps and detachments originate from the same state and that this state arises later in the mechanochemical cycle than the state that gives rise to forward steps. To explain our data, we propose that in each cycle of ATP turnover, forward kinesin steps can only occur before Pi release, whilst backslips and detachments can only occur after Pi release. In the scheme we propose, Pi release gates access to a weak binding K.ADP-K.ADP state that can slip back along the microtubule, re-engage, release ADP and try again to take an ATP-driven forward step. We predict that this rescued detachment pathway is key to maintaining kinesin processivity under load.

**Significance statement:** Kinesin-1 molecular motors are ATP-driven walking machines that typically step forward, towards microtubule plus ends. But they can also step backwards, especially at high load. Backsteps are currently thought to occur by directional reversal of forwards walking. To the contrary, we propose here that kinesin backsteps are not steps, but slips. We show that backwards translocations originate from a different and later state in the kinesin mechanism than the state that generates forward steps. To explain this, we propose that following ATP binding, kinesin molecules that fail to step forward within a load-dependent time window convert to a state that can slip back, rebind to the microtubule, and try again to step forward.

## Introduction

Kinesin-1 is a ubiquitous, ATP-driven molecular motor that moves in 8 nm steps (1) towards the plus ends of cellular microtubules (MTs). To function effectively as an intracellular transporter (2, 3), kinesin needs to move vesicles and other cargo up to several microns in diameter against appreciable hindering loads created by the crowded and dynamic intracellular environment. Understanding how kinesin makes sustained progress under load is an important problem. There is firm evidence that between steps, kinesin waits for ATP with one motor domain (the holdfast head) bound to the MT and the other (the tethered head), poised to step but prevented from doing so until ATP binds (4–10). At low hindering loads, forward steps, towards MT plus ends, predominate, whilst at higher loads, forward stepping slows down and more backsteps are seen. It is clear that both forward steps (11–13) and backsteps (14, 15) require ATP and that at stall, forward steps and backsteps are equally likely (14, 15) – indeed this property defines the stall force. Current models, including our own, envisage that backsteps occur by a hand-over-hand mechanism that resembles that for forward stepping, but with load-dependent reversal of the directional bias (14–19). Nonetheless, it remains possible that forwards and backwards kinesin steps occur by different mechanisms.

To test this point experimentally, we compared the average dwell times for forward steps, backsteps and detachments, for kinesin moving on various types of microtubule. The dwell times are the waiting times preceding each step, consisting of the time spent waiting for ATP to bind plus the time taken to process ATP and complete the coupled mechanical step. We also made counts of forward steps, backsteps (on average 8 nm, but up to 12 nm), larger backsteps (> 12 nm) and full detachments, for kinesin stepping under defined hindering loads between 2 and 9 pN applied with an optical trap. (Fig. 1*A, B*). In a full detachment, the motor releases completely from the MT, causing the bead-motor complex to relax back into the centre of the optical trap (Fig. 1*A*).

**Fig. 1.**
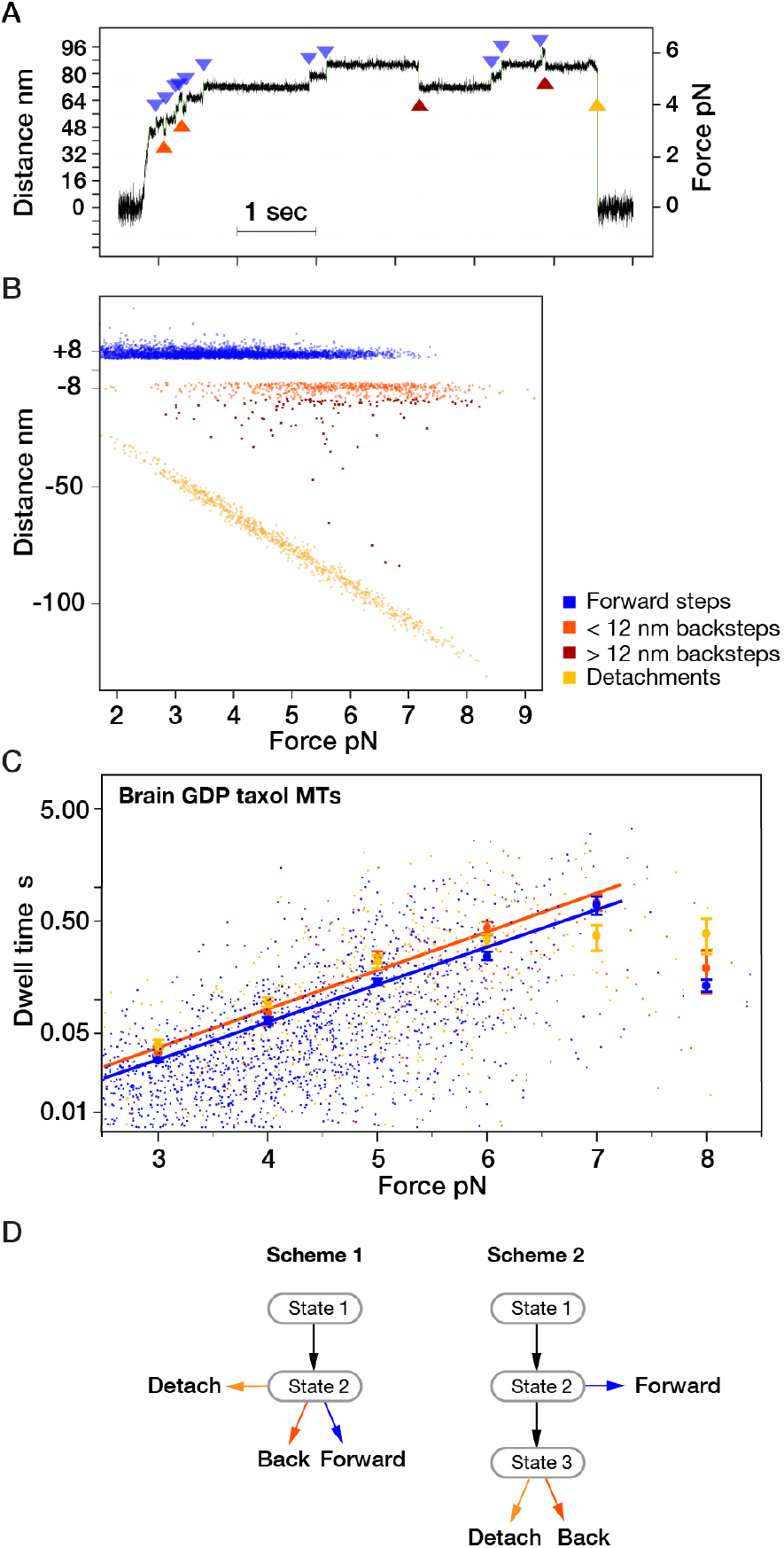
Stepping of single kinesin molecules on brain GDP taxol MTs under load. (*A*) representative sequence of kinesin steps, with examples of each step-class highlighted. (*B*) Step amplitudes and incidence versus hindering force for forward steps, backsteps, larger backsteps and detachments. Trap stiffness for these experiments was ~0.06 pN/nm. (*C*) Forward step (blue), backstep (red) and detachment (yellow) dwell times vs. hindering force for Brain GDP MTs stabilised with taxol. The larger symbols are mean dwell times calculated for bins set at 1 pN intervals. Below stall, dwell times depend exponentially on load, forward step dwell times are characteristically shorter than backstep or detachment dwell times and backstep and detachment dwell times are indistinguishable. Approaching stall force on all types of MT, dwell times reach a plateau, above which further increases in force have little influence on dwell time (15). Mean dwells up to the stall force (7.2 pN) were fit by least squares using log(y) = kx + b. Parameters: blue: k=0.77, b=−5.7 orange: k=0.79, b=−5.6; Errors are SEM. (*D*) Minimal schemes for the kinesin mechanism (see text).

### Dwell times for forward steps are shorter than dwell times for backsteps

For kinesin-1 moving on taxol-stabilised pig brain GDP MTs, the general form of the dwell time versus load relationship is as previously reported (14–17), with forward step dwell times exponentially dependent on load at forces up to and including the stall force (Fig. 1*C*). Crucially however, we find that at substall loads, forward steps have shorter average dwell times than either backsteps or detachments (Fig. 1*C*), whilst backsteps and detachments have similar or identical average dwell times (see next section). Previous work, including our own, has assigned backsteps and forward steps as alternative outcomes from the same state in the kinesin mechanism (14, 15). Correspondingly, current kinesin schemes postulate a minimum of two bound mechanical states (Fig. 1*D*, Scheme 1). In Scheme 1, ATP binds to state 1, the waiting state, and converts it to state 2, the stepping state, which then either steps forward, or backward, or detaches, depending on load. Because in Scheme 1 forward and backward steps originate from the same state, their dwell times are drawn from the same distribution. However we now find that average dwell times for forward steps are shorter than those for backsteps and detachments, whilst those for backsteps and detachments are indistinguishable. To explain this we need an extra state, with ATP binding creating state 2, the try-forward state, and Pi release creating state 3, the backsteps-and-detachments state (Fig. 1*D*, Scheme 2). Scheme 2 is the minimum scheme required to produce different dwell time distributions for forward and backwards steps.

The difference between average forward and backstep dwell times for kinesin moving on GDP taxol brain MTs is just a few milliseconds at 3 pN load, increasing approximately exponentially to several tens of milliseconds at 6 pN load (Fig. 1*C*). On other types of MT lattice, average forward and backstep dwell times are more obviously different. We measured kinesin-1 single molecule stepping mechanics on brain GDP taxol MTs, brain GDP MTs without taxol, brain GMPCPP epothilone MTs and *S. pombe* GMPCPP epothilone MTs. Epothilone is a MT-binding drug that stabilizes both brain and yeast MTs, whereas taxol only stabilizes brain MTs. Previous work has shown that the nucleotide state of tubulin can influence MT sliding velocity in unloaded motility assays (20). Further, lattice expansion and other lattice effects due to changes in nucleotide state (21) drug occupancy (22) or kinesin occupancy (23, 24) can also potentially influence kinesin binding and stepping. Under load, we find that changing MT lattice type indeed substantially affects kinesin stepping mechanics. Most obviously, stalls are shorter on brain MTs (Fig. 2*A*) than on yeast (*S. pombe*) MTs (Fig. 2*C*), indicating that detachment of kinesin-1 at high force is more probable for brain MTs.

**Fig. 2.**
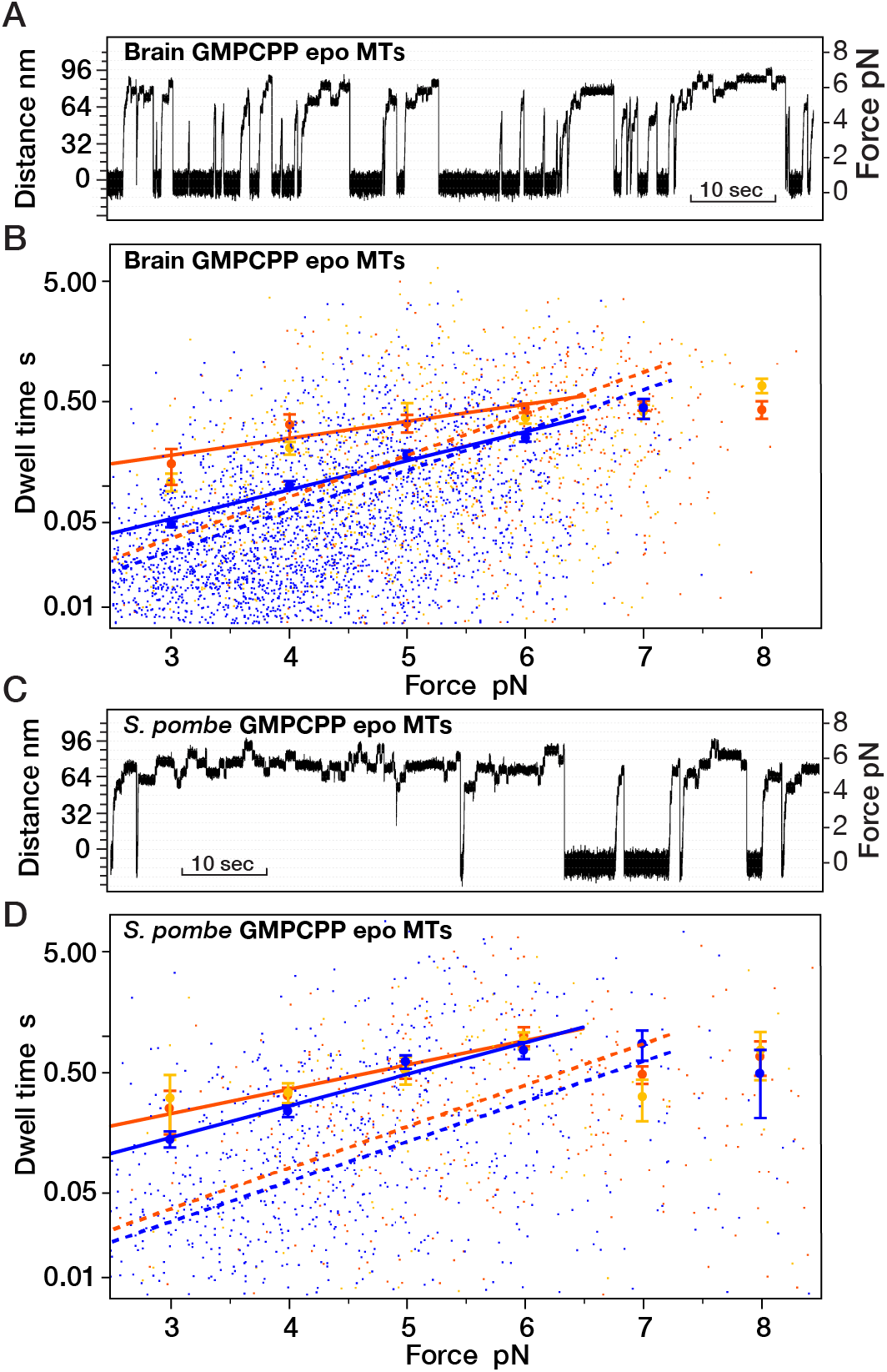
Mechanics of kinesin stepping under load on different MT types. (*A,C*) optical trapping position versus time traces showing the much shorter stalls typically seen with brain GMPCPP epothilone MTs (*A*) compared to *S. pombe* GMPCPP epothilone MTs (*C*). (*B,D*) Dwell times versus load for kinesin stepping on (*B*) Brain GMPCPP epothilone MTs and (*D*) *S. pombe* GMPCPP epothilone MTs. As in Fig 1C, dwell times depend exponentially on load, mean forward step dwell times are characteristically shorter than mean backstep or detachment dwell times and mean backstep and detachment dwell times are indistinguishable. Approaching stall force, backstep dwell times reach a plateau, above which further increases in hindering load have little effect on dwell time (15). Mean dwell times in the region below stall force were fit using log(y) = kx + b. Parameters: (B) blue: k=0.53, b=-4.4; orange: k=0.25, b=-2.3; (D) blue: k=0.6, b=-3.6; orange: k=0.46, b=-2.7; Errors are SEM. In *(B.D)* fits for GDP taxol MTs are shown for comparison (opaque lines).

### Dwell times for backsteps and detachments are the same

For all the types of MT that we tried, we find not only that mean forward step dwell times at substall forces are shorter than those for backsteps and detachments, but also that mean dwell times for 8 nm backsteps, longer backsteps and detachments appear identical (Fig. 1*C*, 2*B*, 2*D*), reflecting that the dwell time distributions for 8 nm backsteps, longer backsteps and detachments superpose and implying that these different types of events originate from the same state in the kinesin mechanism (16, 17). Since backstep and detachment dwell times are longer, on average, than forward step dwell times, this progenitor state must occur later in each cycle of the mechanism than the state that generates forward steps (Fig. 1*D*, Scheme 2).

### Counts of backsteps and detachments vary reciprocally on different MT lattices

Making counts of forward steps, backsteps and detachments at each load gives further insight. The balance between 8 nm backsteps, longer backsteps and detachments at any particular load shifts substantially depending on the type of MT lattice, whereas the fraction of forward steps is little affected (Fig. 3*A-D*). The ratio #forward steps / #backsteps decreases exponentially with load, as previously reported (14, 15), but both the exponential factor and the stall force are different for the different MT lattices (Fig. 3*E*). Stall force is defined as the force at which #forward steps equals #backsteps. For GDP taxol brain MTs, stall force is 7.3±0.2 pN as previously reported (14, 15), whereas for the other types of MT, stall force is 6.4±0.1 pN (Fig. 3*E,* p-value < 0.001 vs GDP taxol brain MTs). By contrast, the ratio #forward steps / {#backsteps + #detachments} versus load is almost invariant for the different types of MT (Fig. 3*F*), reflecting that switching MT lattice type changes the relative incidence of 8 nm backsteps, longer backsteps and detachments, without appreciably changing the count of forward steps or the total count of # backsteps + #detachments at any particular load (Fig. S1). This is only possible in a 3-state minimal scheme (Fig. 1*D*), where backsteps and detachments are alternative outcomes from state 3 and this state succeeds state 2, the state from which forward steps originate. Because detachments and backsteps both originate from state 3, we hypothesized that state 3 is a weak binding state that can diffuse back along the MT and then convert to strong binding to complete a backstep. To test this idea, we varied conditions further.

**Fig. 3.**
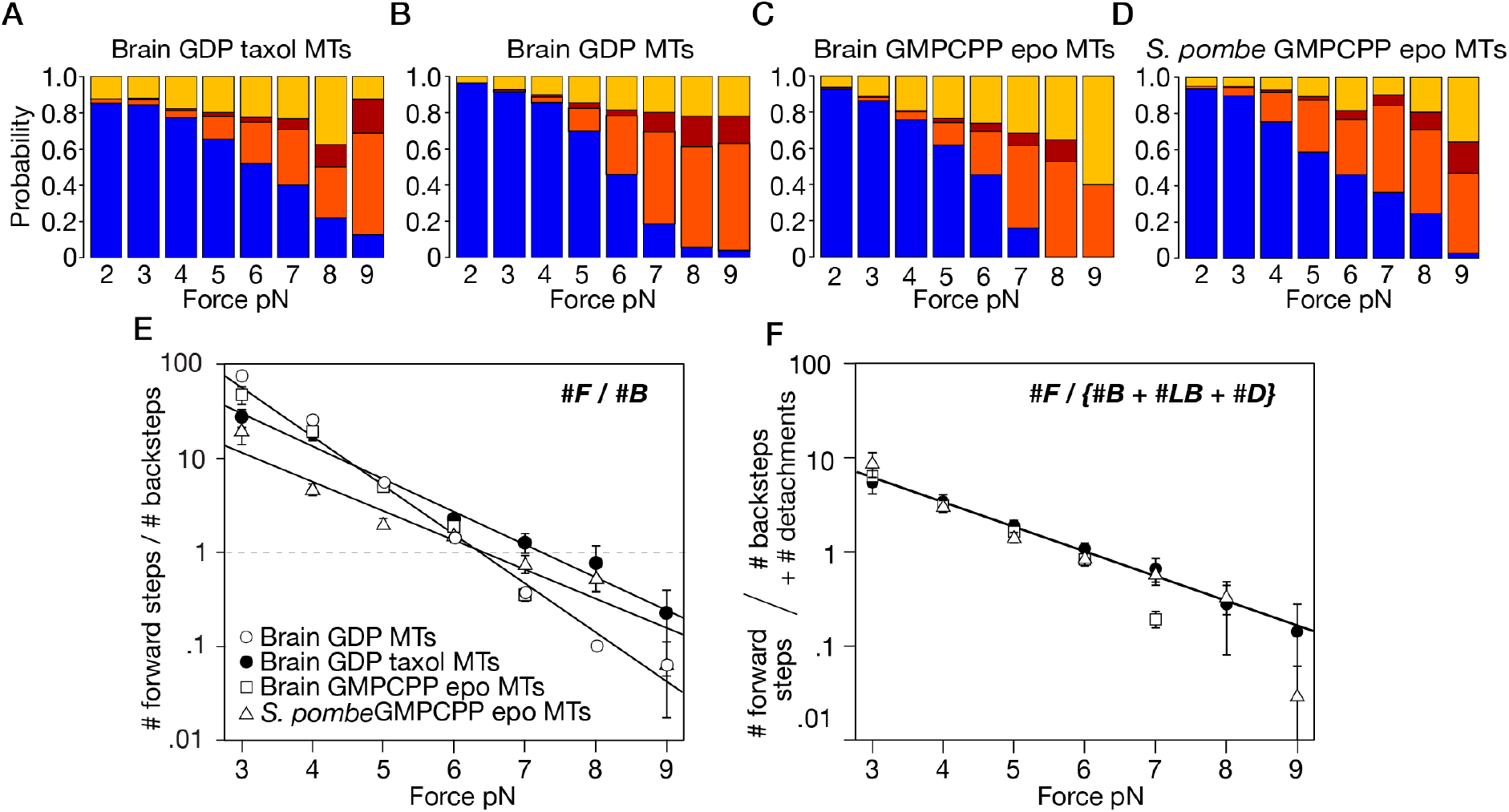
Effects of changing MT type on step-event probabilities under load. (*A-D*) Probability of forward steps (blue), <12 nm backsteps (orange), >12 nm backsteps (magenta) and detachments (yellow) versus force for kinesin stepping on (*A*) Brain GDP taxol MTs, (*B*) Brain GDP MTs (*C*) Brain GMPCPP epothilone MTs (*D*) *S. pombe* GMPCPP epothilone MTs. (*E*) #forward steps / #backsteps vs. force, for kinesin stepping on these four different MT types. Data were fit to log(y) = kx + b, weighting the fit by the reciprocal SE. For brain GMPCPP epothilone MTs k=-1.16, b=7.45; for *S. pombe* GMPCPP epothilone MTs k=-0.79, b=5.0, for brain GDP taxol MTs: k=-0.83, b=5.9. Errors are SE. (*F*) #forward steps / {#backsteps + #larger backsteps + #detachments} versus force, for kinesin stepping on different MT types. Fit shown is to brain GDP taxol MTs data: k=-0.58, b=3.5.

### Added ADP inhibits forward steps and promotes backsteps and detachments

K.ADP is the weakest state of the kinesin cycle (25, 26). If, as we hypothesise, state 3 is a K.ADP state, then adding extra ADP to the bathing solution should enrich state 3 via ADP binding to the waiting state. To test this possibility, we recorded kinesin stepping on brain GDP taxol MTs in the presence of added ADP (Fig. 4*B*). Increasing [ADP] without changing [ATP] favours backstepping and disfavours forward stepping (Fig. 4*B,F*), thereby reducing stall force from 7.3±0.2 to 5.1±0.2 pN (Fig. 4*E*). More detailed counts show that added ADP not only produces more backsteps, but also adds a population of longer backsteps (Fig. S2). These data indicate that adding ADP enriches state 3, from which backsteps and detachments arise (Fig. 1*D*), consistent with state 3 being a K.ADP-K.ADP state and with added ADP enriching the K.ADP-K.ADP state by binding to the waiting state (the K – K.ADP state). Dwell time distributions (Fig. 5) show that added ADP increases the average forward step dwell time at low load and abrogates forward stepping at high load, consistent with the main element of dwell time under load being the waiting time before nucleotide binding and with added ADP competing directly with ATP for binding to the waiting state. The change in the dwell time distribution caused by adding ADP suggests that some events might involving multiple rounds of ADP release and rebinding.

**Fig. 4.**
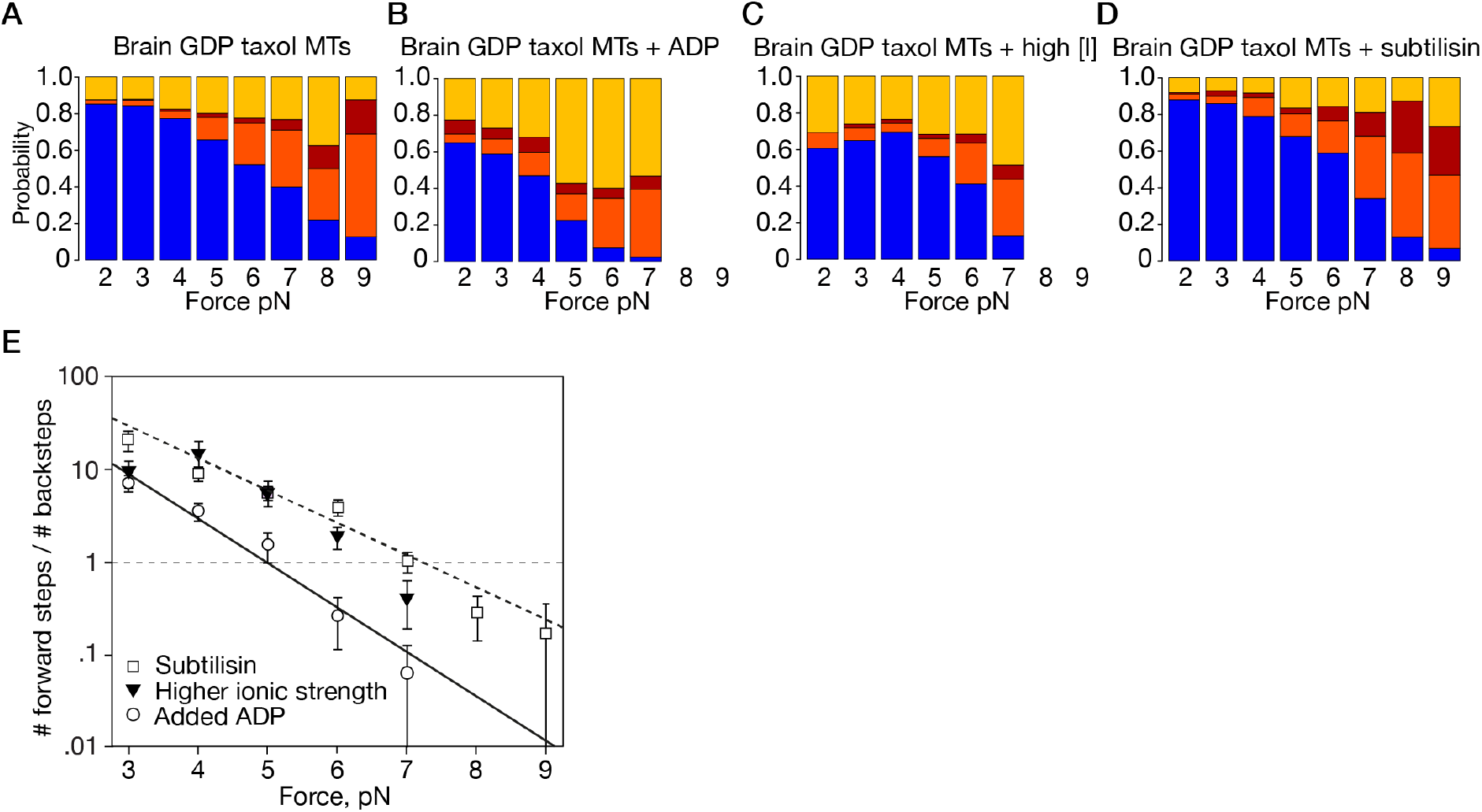
Effects of changing conditions on step-event probabilities under load. (*A*) Brain GDP taxol MTs, as in Fig. 3*A*. (*B*) Brain GDP taxol MTs with added 1mM ADP. (*C*) Brain GDP taxol MTs at higher ionic strength (BRB160 instead of BRB80) (*D*) Brain GDP taxol MTs treated with subtilisin (*E*) Effect of ADP, higher ionic strength and subtilisin treatment on #forward steps / #backsteps ratio, for kinesin stepping on GDP-taxol MTs under hindering load. ADP data fit parameters: k=-0.69, b=4.9. Broken black line is the fit for stepping on GDP taxol MTs, from Fig. 3E. Errors are SE.

**Fig. 5.**
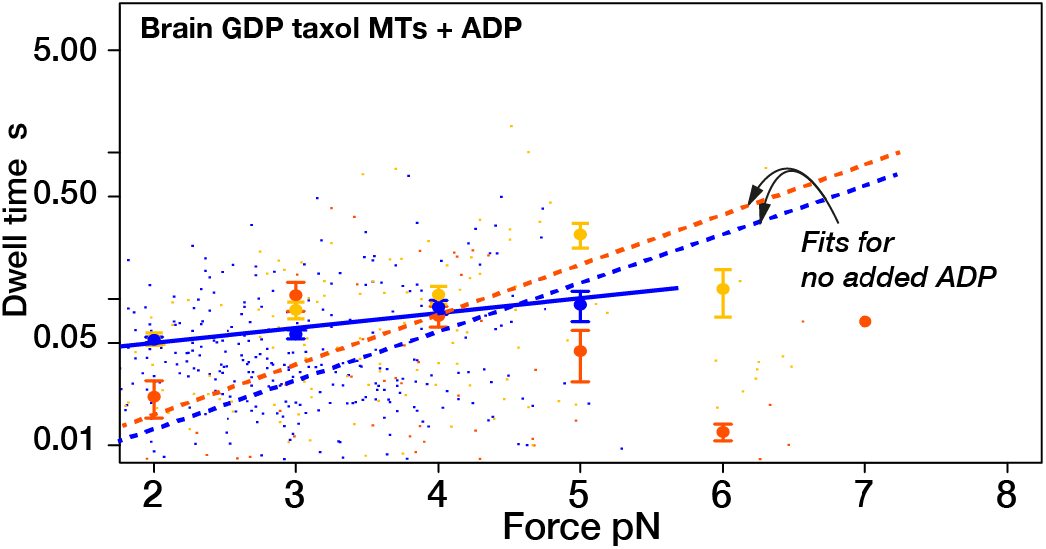
Dwell time distributions with added ADP. Compared to stepping on GDP taxol MTs with no added ADP (Fig 1*B* fits shown here for reference as dotted lines), adding ADP reduces the load dependence of the mean dwell time for forward steps.

### Increased ionic strength promotes detachment

By increasing ionic strength, we expect to weaken our hypothesized weak binding K.ADP-K.ADP state 3, because weak binding has a considerable electrostatic component (27). We find that indeed increased ionic strength promotes detachment, with much smaller effects on ~8 nm backsteps and longer backsteps (Fig. 4*C*), consistent with increased ionic strength weakening and depopulating the K.ADP-K.ADP state, without appreciably affecting forward stepping. As a result, stall force is only marginally reduced (Fig. 4*E*).

### Subtilisin treatment of MTs lengthens backsteps and inhibits detachment

Cleaving the C-terminal E-hook peptides of tubulin (28) has only slight effects on kinesin velocity at low load, but reduces processivity (29). To further probe the nature of our envisaged semi-detached state, we cleaved the E-hooks from GDP-taxol MTs using subtilisin, and again examined stepping mechanics under load. Subtilisin digestion has little or no effect on the tendency to take forward steps relative to all other events (Fig. 4*D*). Subtilisin digestion does however increase the probability of long (>12 nm) backsteps and decrease the probability of detachments. These data again demonstrate that the balance between short backsteps, long backsteps and detachments can be shifted substantially by experiment, without affecting forward stepping, consistent with our contention that forward steps originate from a different and earlier state in the kinesin mechanism than the state generating backsteps and detachments.

### Proposed Model

Our key finding is that dwell times for kinesin backsteps, long backsteps and detachments are drawn from the same dwell time distribution, whilst the dwell times for forward steps are drawn from a different distribution with a shorter mean dwell time. Scheme 2 (Fig. 1*D*) is the minimum scheme necessary to explain these findings. Scheme 2 is needed to explain how shifting kinesin between different MT types can shift the balance between backsteps and detachments, without affecting forward stepping. Scheme 1, in which forward steps and backsteps originate from the same state, is ruled out, because in scheme 1 forward steps and backsteps originate from the same state and so should originate from the same dwell time distribution – and we have shown that they do not.

How might scheme 2 map to the kinetic mechanism of kinesin? In our proposed new model (Fig. 6), ATP binding opens a time window devoted exclusively to forward stepping. Within this time window, our scheme allows for forward stepping under load from either the ATP or the ADP.Pi states of the holdfast head (blue pathway). We include both these possibilities because whilst recent evidence points to the K.ADP.Pi - K.ADP state as the origin of forward steps at zero load (10), the effects of load on the hydrolysis step are not yet clear. It is possible, for example, that the hydrolysis step might be reversible under load, as it is for myosin (30, 31). In our model, the forward-step time window is closed by Pi release (32) and we propose that backsteps and detachments occur only after Pi release. We envisage that at high hindering loads, forward stepping increasingly fails to occur within its time window and that closure of this time window by Pi release switches the motor into a weakly bound K.ADP - K.ADP state (orange pathway). In our scheme, this K.ADP - K.ADP state then typically slips back along the lattice, re-engages, releases ADP and returns to its K – K.ADP waiting state, completing a *de facto* backstep. Under backwards load, re-engagement typically occurs to the closest available tubulin heterodimer, resulting in an 8 nm backslip. Less routinely, backslips of 16 or 24 nm can occur, or the motor can fully detach. For completeness, we include a presumptively off-pathway transition, in which MT-activated ADP release occurs with the motor standing in place (grey arrows). It is unclear at present to what extent this pathway is used. It is also unclear whether our K.ADP - K.ADP slip state rebinds to the same protofilament (PF) from which it unbound, or to a different PF. We also do not yet know whether following a slip, the same head that detached then re-engages, or whether the two heads can swap over. Distributions of backstep sizes may provide a clue. Counts of backsteps in each size-class reveal approximately exponentially decreasing likelihoods of 8, 16, 24 and 32 nm backsteps under load, for each of the MT lattice types that we tested (Fig. 7*A-D*). This suggests a fixed probability of rebinding (rescue) of our hypothesised K.ADP-K.ADP slip state at each subunit position back along a single PF, but further work will be needed to clarify this point.

**Fig. 6.**
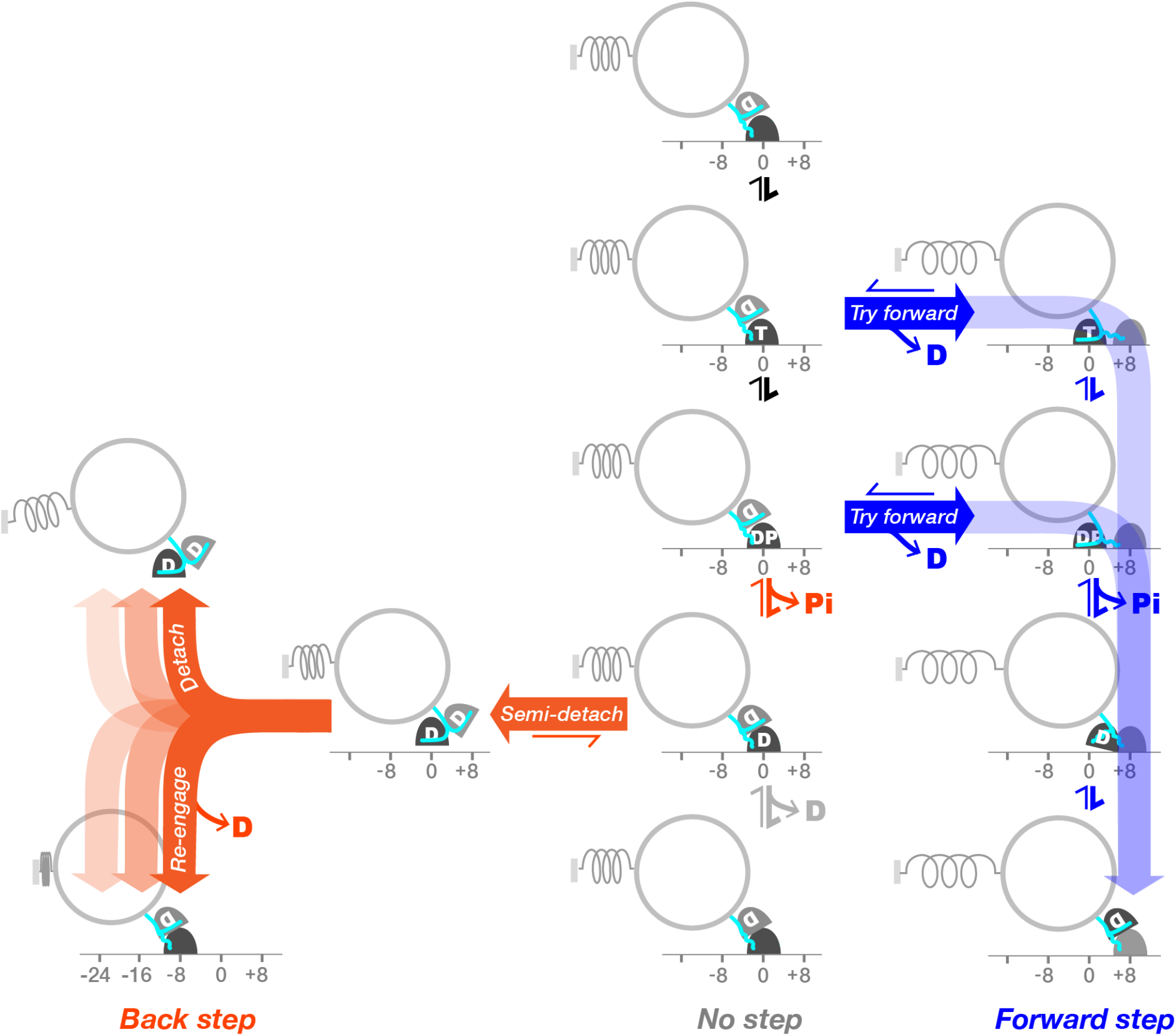
Proposed new model for kinesin stepping under load. The key feature of the model is that forward steps and backward steps originate from different states. During each cycle of ATP turnover under load, kinesin first attempts to step forward from its K.ATP-K.ADP and K.ADP.Pi-K.ADP states (blue arrows). In our model, backsteps do not occur from either of these states. At higher loads, forward stepping increasingly fails to complete before Pi release and kinesin then enters a weak binding K.ADP-K.ADP state that can slip back along the lattice (orange arrows), re-engage, release ADP and begin a fresh attempt to step forward. For completeness, a presumptively minor process, ADP release whilst standing still (grey arrows) is also shown.

**Fig.7.**
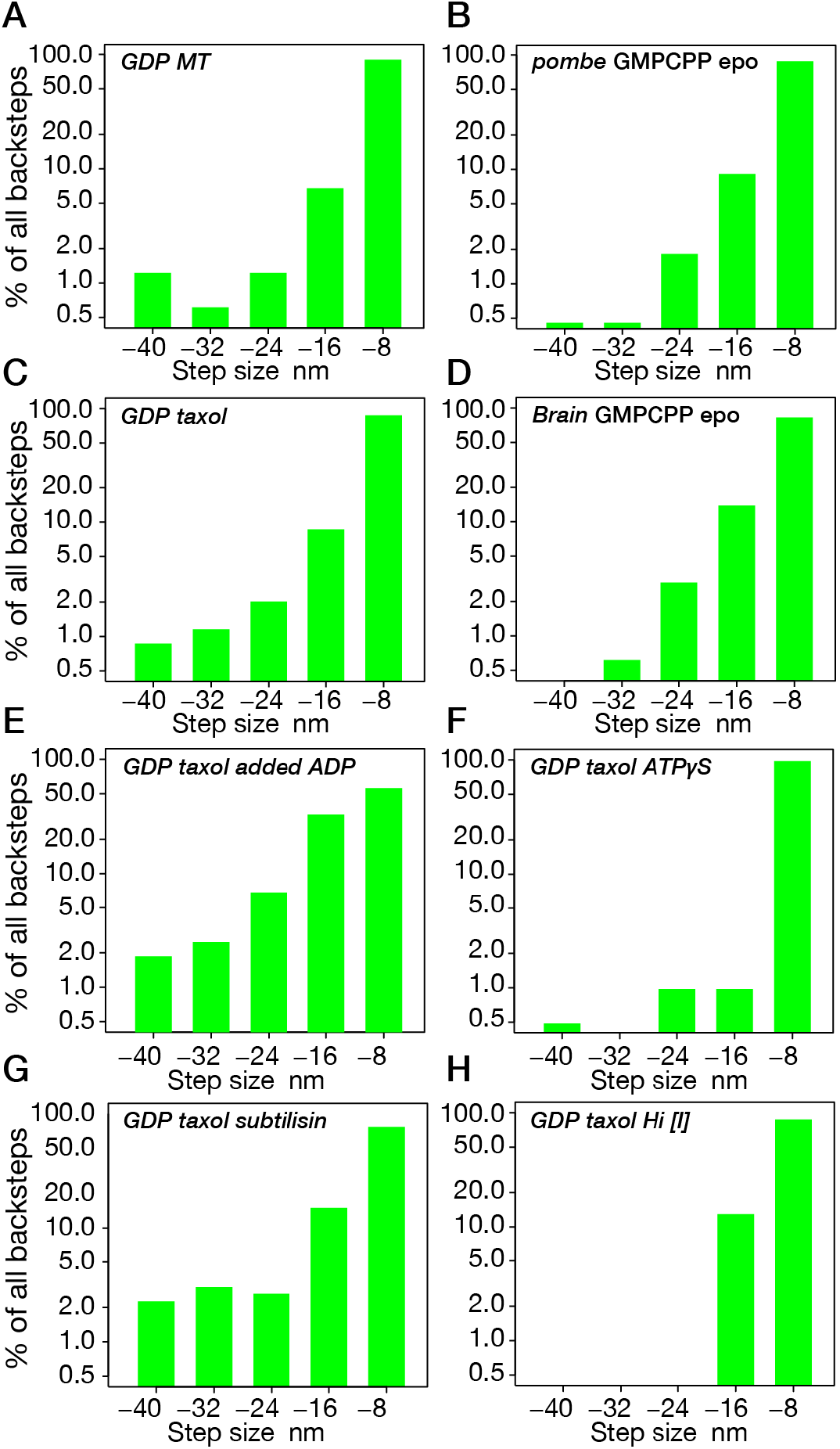
Backstep size distributions. (*A-H*) percentage of all backwards events in each size class, for different MT lattices and in different conditions. Note Y axis is on a log scale. (*A-D*) Different MT lattices. Probability distributions appear roughly exponential, consistent with backsteps being backslips that have an equal probability of reattachment at each position back along a single PF towards the MT minus end. (E-F) Under perturbation, this exponential relationship no longer holds. In (*E,G*) longer backsteps appear more probable; in (F,H) longer events are very rare.

### Relationship to other models

Our new model (Fig. 6) is conventional in that ATP binding gates forward stepping, with gating involving a combination of ATP-dependent unparking of the leading (tethered) head (4, 5, 33, 34) and ATP-dependent stabilization of neck linker docking on the MT-attached trail head (35–38). Assigning K.ADP as the weakest binding state in the cycle is also conventional (25, 26) and our assignment of the K.ADP-K.ADP state as the progenitor for detachments is also consistent with earlier models (10, 15, 39). The idea that some backwards events are slips is also not novel; 16 nm, 24 nm and larger backwards displacements have previously been assigned as slips, on the basis that they are too fast to represent a sequence of 8 nm steps that are each coupled to a full cycle of ATP turnover (15). The novel aspects of our proposed scheme are firstly, that the interlude between ATP binding and Pi release (the hydrolysis time) is devoted *exclusively* to forward stepping, secondly that *all* backwards events at substall loads are backslips and thirdly that backslips and detachments arise from the same K.ADP-K.ADP progenitor state.

Our proposed scheme is also consistent with recent work showing that the rate of kinesin stepping can be influenced either by supplying extra ADP to inhibit tethered head attachment or by lowering ionic strength to strengthen the binding of the K.ADP state to the MT (10). At zero external load, backsteps are rarely seen, but our data suggest that similar effects may operate – that is, that increased [ADP] and ionic strength respectively enrich or deplete a weak-bound K.ADP - K.ADP slip-state (40).

Slip and re-engage behavior under backwards load has been seen before in kinesins. We previously assigned 16nm and larger backwards kinesin-1 events as slips (15). Jannasch et al (41) showed that backslipping of Kip3, the *S. cerevisae* kinesin-8 limits its stall force to < 1 pN. The claim we make here is that 8 nm kinesin-1 backsteps are also slips, so that a slip-and-re-engage pathway is integral to the mechanism of a transport kinesin that can do appreciable mechanical work.

Our proposed scheme is consistent with recent work showing that adding Pi extends the single molecule run length of kinesin-1 (32) under forwards load, because it predicts that adding Pi will enrich the K.ADP.Pi-K.ADP state and so delay the formation of the backslipping K.ADP-K.ADP state. Equally, our scheme is consistent with models in which slower ATP hydrolysis promotes forward stepping by promoting diffusion-to-capture by the tethered head (10, 42), because slower ATP hydrolysis would allow more time for forward stepping. We attempted to test this last point directly using ATPγS, an ATP analogue that drives MT sliding assays at ~2-3% of ATP velocity (43). Using iSCAT, ATPγS was recently shown to support slow, erratic stepping of single kinesin-1 molecules at zero load (44). Under load in the optical trap, we find that ATPγS also drives slow and erratic stepping of kinesin (Fig. 8*A*). However, we find that the MT-activated rate of ATPγS turnover by kinesin in solution is only 6-7 fold lower than that for ATP (Fig. 8*D*). These data suggest that ATPγS is hydrolysed relatively rapidly, but has poor mechanochemical coupling, so that at present it does not provide a useful test of our scheme.

**Fig. 8.**
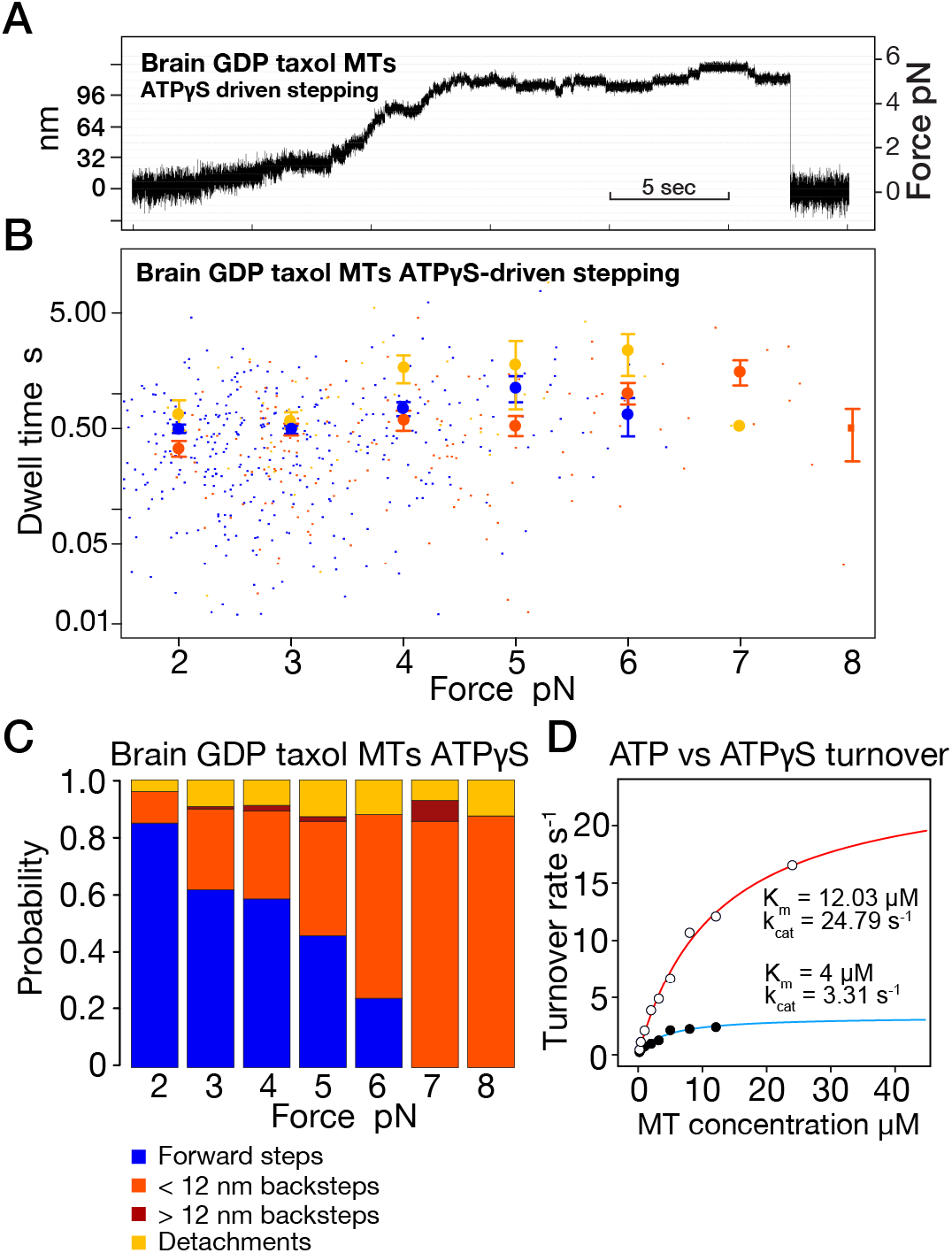
ATPγS – driven stepping. (*A*) Representative optical trapping record. Stepping is slow and steps are less well defined than in ATP, especially at low loads. (*B*) Dwell time versus load plot for forward steps, backsteps and detachments. Dwell times appear less load-dependent than in ATP. (*C*) Event probabilities versus load. Backsteps appear much more probable than in ATP (compare Fig. 3*A*). (*D*) Nucleotide turnover for ATP (red fit) versus ATPγS (blue fit).

### Implications

Most immediately, our proposed new model (Fig. 6) has implications for the mechanochemical coupling of kinesin. As with previous models, in our new model kinesin consumes more ATP at high loads, because backslips consume ATP. In our scheme, even a single 8 nm backslip will consume one ATP to slip back 8 nm and another to step forward to regain its original position. Longer backslips will require correspondingly more ATP-driven forward steps to regain the ground lost. In one sense, these backslips are futile, because they require ATP turnover and generate negative progress. In another sense however, they are far from futile, because they create the opportunity to retry forward stepping under load without letting go of the MT and losing all the ground previously gained. A further possible pathway for futile cycling is denoted by the grey arrow in Fig. 6, corresponding to MT-activated ADP release without stepping. At present we have no information on the flux through this pathway, which would also consume ATP, but without losing ground.

Whilst our data argue that the large majority of kinesin-1 backward translocations are slips, we cannot exclude that a small fraction of true mechanical “hand-over-hand” backsteps, as opposed to backslips, is present. However, since these events would originate from the same state that generates forward steps, they would draw from the same dwell time distribution as forward steps (Fig.1*D*, scheme 1) and tend to make the dwell time distribution for short backsteps more like that for forward steps - whereas we see that the dwell time distribution for short backsteps is different from that for forward steps, and indistinguishable from that for long backsteps & detachments. On this basis we can firmly conclude that at least the great majority of backsteps are slips.

By rescuing kinesins that have failed to step forward within a load-dependent time window and allowing them to retry forward stepping, our proposed rescued detachment pathway increases the ultimate success rate for processive forward stepping under load, with a corresponding increase in stall force and the ability to do useful mechanical work. These gains come at the expense of extra ATP consumption, so that the ATP cost of each net forward step increases substantially at high loads. Effectively, backslipping allows kinesin-1 to change gear under load, by adaptively combining tight-coupled forward steps with loose-coupled backslips.

## Materials and Methods

### Kinesin-beads

Unmodified 560 nm diameter polystyrene beads (Polysciences) were mixed with purified recombinant full-length *Drosophila* kinesin-1 (12), serially diluted until only 1/3 of beads were motile. Experiments were performed in BRB80 buffer (80 mM K-PIPES (pH 7.0), 2 mM MgSO_4_, 1 mM EGTA, 3 g/L glucose) with 1 mM ATP and glucose-catalase oxygen scavenging system and 10 μM taxol or 10 μM epothilone or with no drug.

### Microtubules

Purified tubulin (either from porcine brain or from *S. pombe (α1, α2, β* isoforms (45)) was polymerized in BRB80 at 37°C for 45-60 min in the presence of 1 mM GTP and 2 mM MgCl_2_. MTs were spun down and resuspended in BRB80 supplemented with taxol (Sigma-Aldrich) or epothilone (VWR International) to 20 μM. Since taxol is not effective on *S. pombe* MTs, epothilone was used instead in combination with GMPCPP. In the case of unstabilised GDP MTs, no drug was added but MTs were capped immediately following assembly using tubulin premixed with 1 mM GMPCPP and incubated for 1 h, added to a final concentration of 1 μM. The capped MTs were spun down and resuspended in BRB80.

### Flow cells

The flow cell surface was passivated with 0.1 mg/ml PLL-PEG and then with 0.1 mg/ml casein (Susos AG, Dübendorf, Switzerland). MTs were covalently attached to the coverslip surface using mild glutaraldehyde crosslinking to the APTES silanized surface (46).

### Optical trapping

A custom-built optical trap (15) was used, equipped with a 6 W Nd:YAG 1064 nm laser (I E Optomech Ltd, Newnham, England). High ionic strength experiments were performed in BRB160, which is identical to BRB80 but with 160 mM K-PIPES. MTs were initially visualized by differential interference contrast (DIC) microscopy and beads moved into position above the MTs by steering the trap. Imaging was then switched to amplitude contrast and the image projected on to the quadrant photodiode detector. Data were recorded at 20 kHz and moving-average filtered to 1 kHz during analysis.

### Data analysis

Data were analysed in R using custom-written code (available on request). Automated step-detection was implemented using t-test analysis. In the t-test analysis, 8 data points before the suspected step and 8 after the step were compared by t-test. The threshold t-value was set to 30 and the minimum step size to 5 nm. Dwell times were defined as the waiting time between two consecutive steps. ~30-40% of the longest dwells in each force bin were manually verified to ensure there were no undetected steps within these dwells. Only steps above 2 pN could reliably be detected and only these were processed. Below 3 pN backsteps were rare and above 8 pN forward steps were rare. For forward step:backstep ratio measurements, the force range 3-8 pN was analysed to ensure a sufficient number of both forward steps and backsteps. Each force bin includes data at the force shown ± 0.5 pN.

## Acknowledgements

We thank Douglas Martin, Masanori Mishima, Douglas Drummond and Huong Thuy Vu for helpful comments.

## Funding

This work was funded by a Wellcome Senior Investigator award to RAC, grant number 103895/Z/14/Z.

## Author contributions

AT performed all the experiments and analysed all the data. NC designed and built the optical trap. AT, NC and RC collaborated to design and interpret experiments and write the ms.

## Competing interests

Authors declare no competing interests.

## Data and materials availability

All data will be made available.

## Supplementary Figures

**Supplementary Fig. 1.**
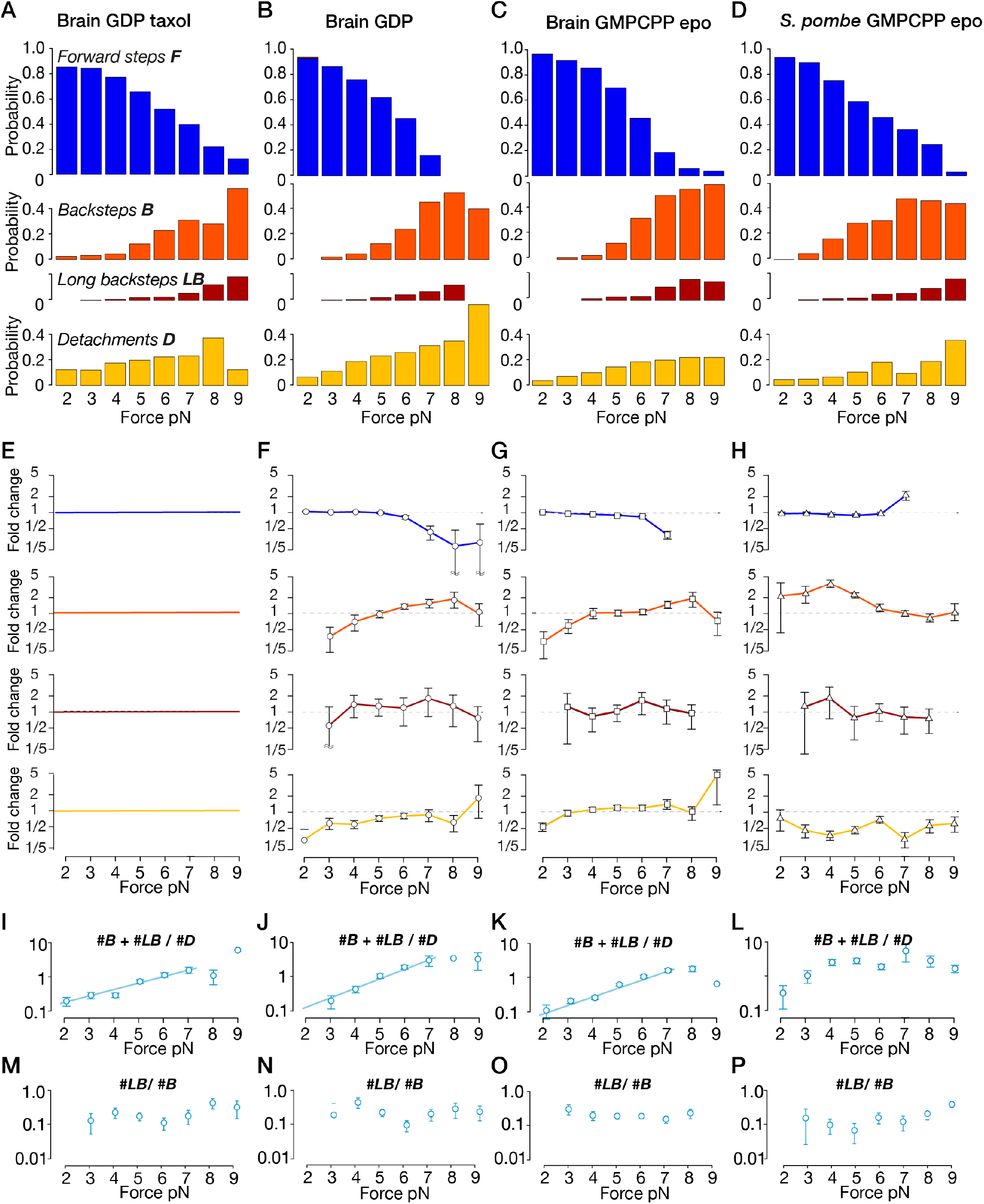
Event counts versus MT lattice type. Further analysis of relative event counts versus load. (*A-D*) Relative event counts for different lattice types; replots of the data shown in Fig.3*A-D*, facilitating comparison of the effects of changing MT lattice type. (*E-H*) fold change in these same event counts, compared to GDP taxol MTs. (*F*) and (*G*) are relative to brain GDP taxol MTs; (*H*) is relative to brain GMPCPP epothilone MTs, so that only the species, and not the species and the drug, varies. (*I-L*) ratio of all backstep events to detachments versus load, for different lattice types. (*M-P*) ratio of long backsteps to 8 nm backsteps versus load, for different lattice types.

**Supplementary Fig. 2.**
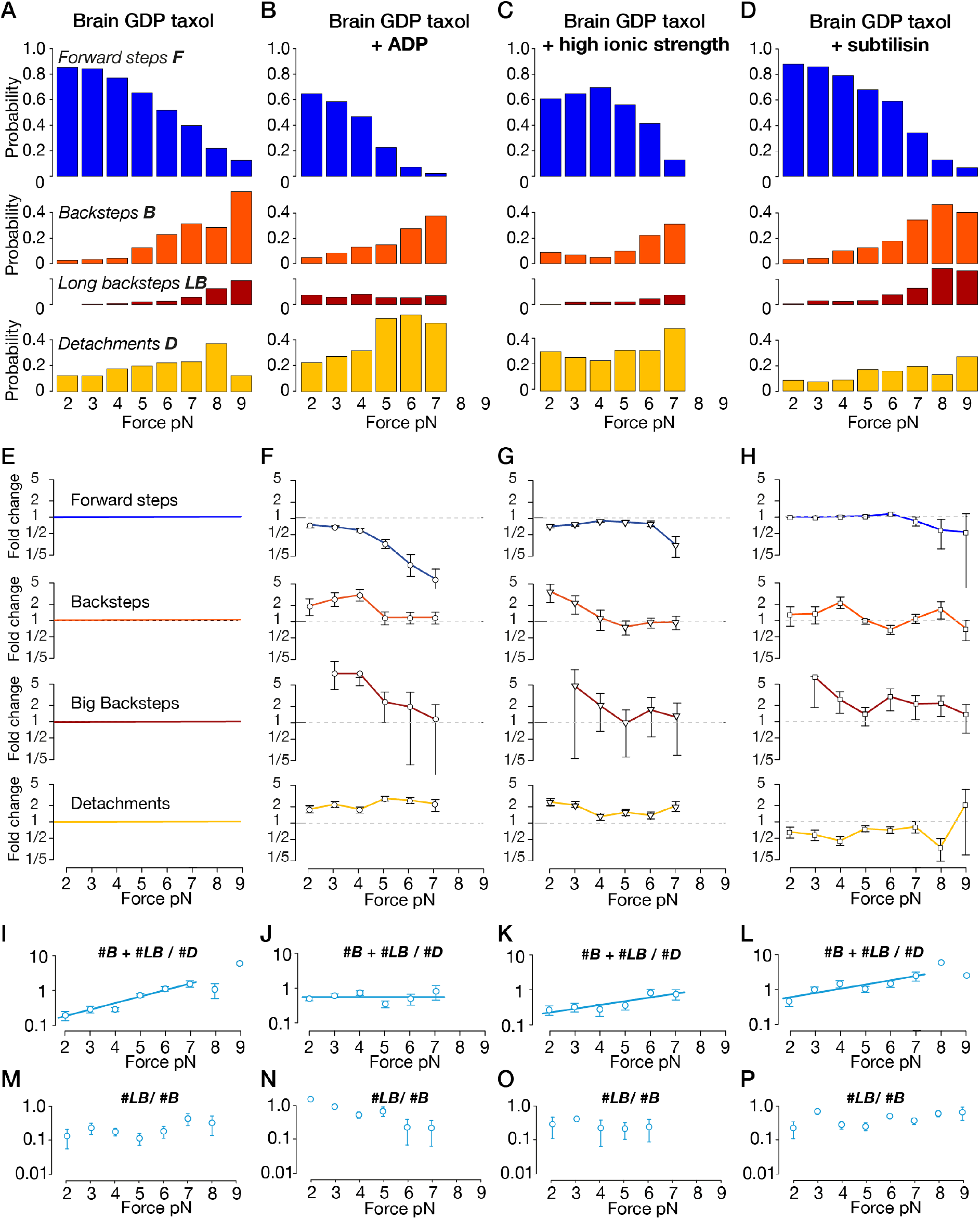
Event counts under varying conditions. Further analysis of relative event counts versus load. (*A-D*) Relative event counts for different conditions, replots of the data shown in Fig.4*A-D*, facilitating comparison of the effects of changing MT lattice type. (*E-H*) fold change in these same event counts, compared to GDP taxol MTs. ADP suppresses forward stepping, ehnaces backstepping and detachment, and adds a population of long backteps at low load that reduces with increasing load. (*I-L*) ratio of all backstep events to detachments versus load, for different conditions. (*M-P*) ratio of long backsteps to 8nm backsteps versus load, for different lattice types.

## Notes

### Competing Interest Statement

The authors have declared no competing interest.

### Summary of Updates

Revision 2 in response to further reviewer comments.

